# ACR TI-RADS^®^ SCORE ULTRASOUND – PICTORIAL ESSAY OF ULTRASOUND / ELASTOSONOGRAPHY / ANATOMY / CYTOLOGY and HISTOLOGY

**DOI:** 10.1101/2022.11.29.518434

**Authors:** Luís Jesuíno De Oliveira Andrade, Luis Matos De Oliveira, Alcina Maria Vinhaes Bittencourt, Letícia Góes De Carvalho Lourenço, Gabriela Correia Matos De Oliveira

## Abstract

**Introduction:** Thyroid Imaging Reporting and Data System (TI-RADS^®^) is a system for classifying thyroid nodules detected at ultrasonography, aiming at a descriptive standardization as well as to classify their risk of malignancy based on sonographic findings.

**Aium:** To present a pictorial essay by composing anatomical, histological, ultrasound and elastography images of the TI-RADS^®^ score.

**Method:** Using software for image composition, based on the ultrasound image of the various types of thyroid nodules, we adapt to the anatomical image, the elastosonography, and histological corresponding TI-RADS^®^ and present a pictorial essay.

**Results:** The correlation between the sonographic, elastography, anatomical, cytology and histological images corresponding to the TI-RADS^®^ score are demonstrated.

**Conclusion:** Ultrasonographic features, elastosonography, anatomical, and histological in evaluation of thyroid nodules can correlate with features TI-RADS® score.

## INTRODUCTION

Thyroid nodules are very prevalent in the adult population; however, about 90% are benign. Ultrasound is the imaging method usually employed in the analysis of thyroid nodules. However, ultrasound findings are in many occasions non-specific, and the conclusive diagnosis is defined by fine needle aspiration biopsy or even by histological examination after surgery. Multiple echographic studies have been proposed to characterize the risk of malignancy of the nodules (1).

Horvath et al. (2) based on the Breast Imaging Reporting and Data System (BI-RADS) model of American College of Radiology (ACR) (3) decided to evaluate the possibility of applying the BI-RADS in the sonographic evaluation of thyroid nodules, grouping the nodular characteristics into different categories with a percentage of malignancy analogous to the BI-RADS, which she called Thyroid Imaging Reporting and Data System (TIRADS).

In order to provide physicians with evidence-based suggestions for therapeutic management based on a grouping of well-established sonographic patterns or expressions that can be adopted for each lesion, the ACR organized a committee and developed TIRADS, standardizing the diagnostic analysis of thyroid nodules through the elaboration of a lexicon (4).

As per the ACR-TIRADS system (Figure 1), five characteristics of the nodule are included in the scores, including (1) component (choose one): cystic or almost completely cystic 0; spongiform 0; mixed cystic and solid 1; solid or almost completely solid 2; (2) echogenicity (choose one): anechoic 0; hyperechoic or isoechoic 1; hypoechoic 2; very hypoechoic 3; (3) shape (choose one): wider-than-tall, 0; taller-than-wide, 3; (4) margin (choose one): smooth 0; Ill-defined 0; lobulated or irregular 2; extrathyroid extension 3; and (5) echogenic foci (choose one): none or large comet-tail artifact 0; macrocalcification 1; peripheral (rim) calcification 2; punctate echogenic foci Thus, the echographic aspects of nodules in the ACR-TIRADS are scored according to their characteristics, and classified as benign (TIRADS 1), not suspicious (TIRADS 2) minimally suspicious (TIRADS 3), moderately suspicious (TIRADS 4), highly suspicious for malignancy (TIRADS 5). The indications for follow-up with ultrasound or fine needle aspiration cytology are based on the ACR-TIRADS score of the nodule as well as its maximum diameter (5). Sonoelastography because it is not yet available on many ultrasound machines, as well as autoimmune thyroiditis due to its scarcity have not yet been added categorically to the ACR-TIRADS classification.

**Figure 1.**
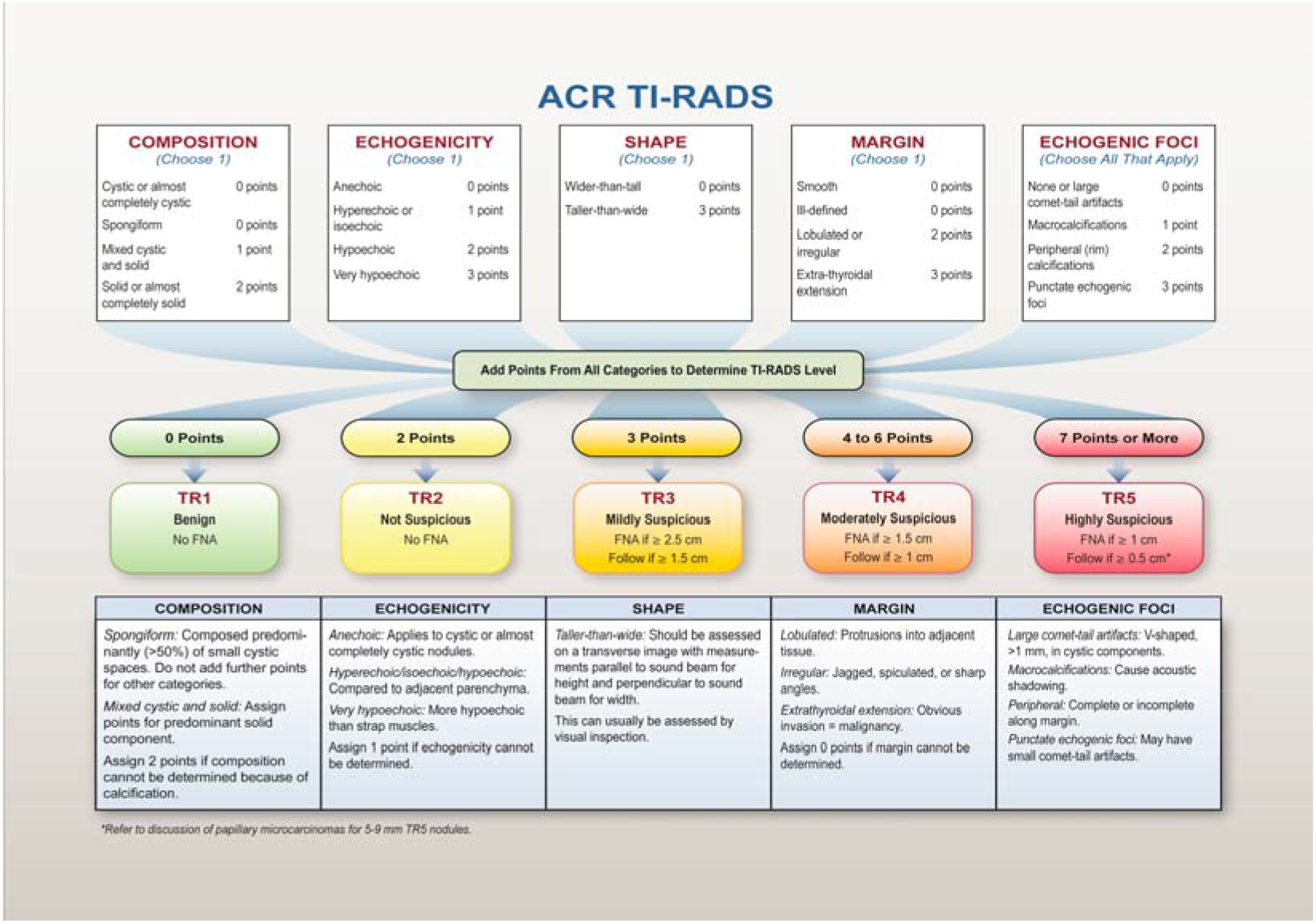
ACR-TIRADS classification. **Source**: Journal of the American College of Radiology. Volume 14 Issue 5 Pages 587-595 (May 2017) DOI: 10.1016/j.jacr.2017.01.046

Elastosonography assists as a non-invasive ultrasound method that evaluates tissue strain. Lately it has been widely used in the evaluation of thyroid nodule. In such a way that both shear wave elastography as for real-time elastography show excellent results in assessing the risk of malignancy show excellent results in assessing the risk of malignancy, using nodule hardness as a criterion of suspicion for malignancy (6). In strain elastography a red-green-blue color map is displayed: red it has been agreed that characterizes soft tissue, green represents intermediate hardness (equal strain), and blue characterizes hard nodules (no strain) (7). Thus, scores have been proposed to assess tissue elasticity, such as the Asteria et al. (8) score, in which score 1 corresponds to a totally green nodule; score 2 to a predominantly green nodule, with some blue areas; score 3 to a predominantly blue nodule, with some green areas; score 4 to a totally blue nodule (Figure 2).

**Figure 2.**
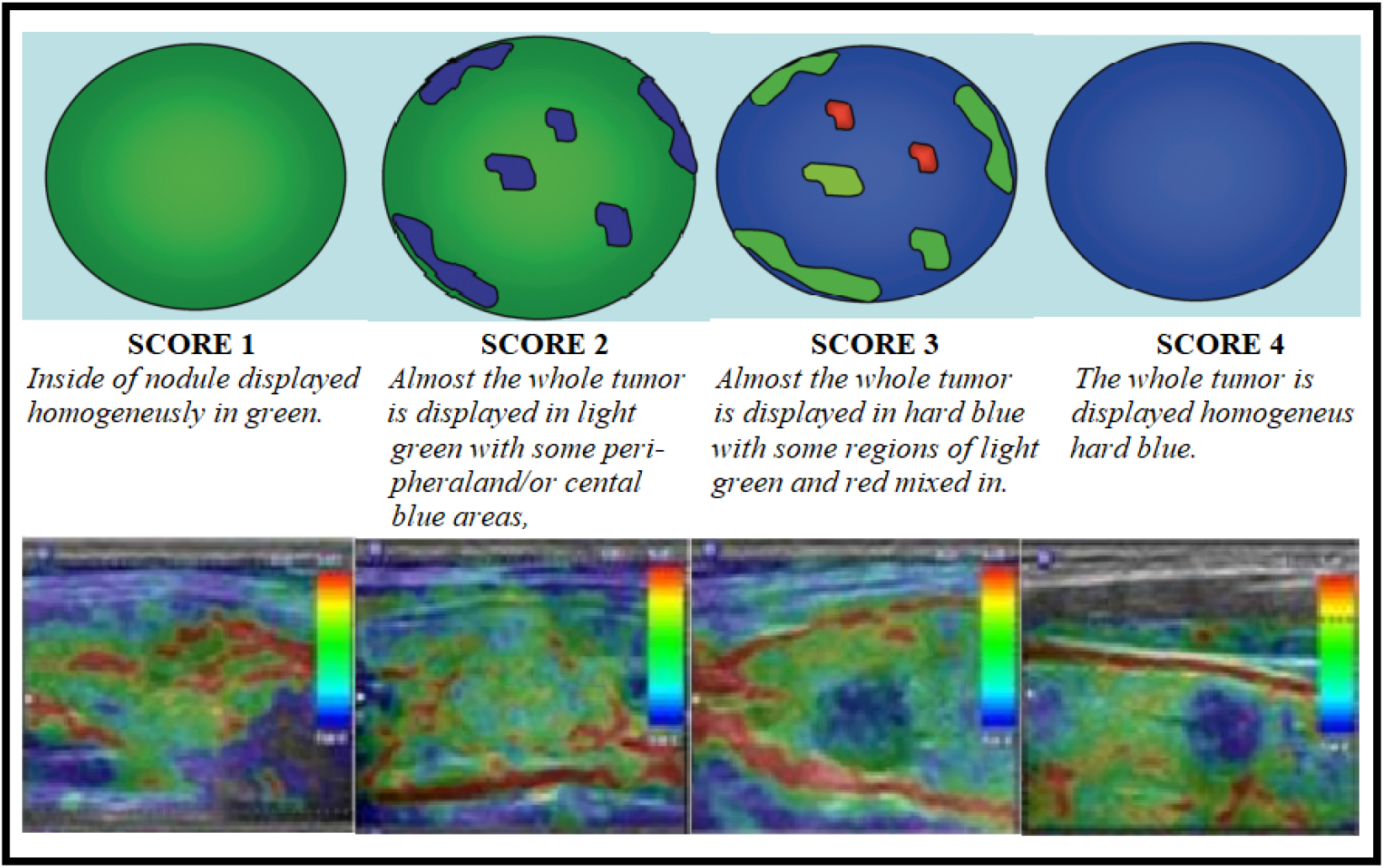
Qualitative evaluation of elastography Strain. **Source**: Asteria C, Giovanardi A, Pizzocaro A, Cozzaglio L, Morabito A, Somalvico F, et al. US-elastography in the differential diagnosis of benign and malignant thyroid nodules. Thyroid. 2008;18(5):523-31.

Scores 1 and 2 are considered benign and scores 3 and 4 are suspicious for malignancy (9). In addition to the color score, tissue elasticity can be assessed using the strain ratio, a measure indicated by the device, which compares the elasticity of the thyroid nodule and the adjacent thyroid tissue (9).

Although numerous imaging examinations have been used in day to day clinical practice, incorrect analysis can occur. This manuscript aims to present a pictorial essay, composing ultrasound images with elastosonography image, anatomical image, cytological image and histological image based on ACR-TIRADS score of thyroid nodules.

## METHOD

Based on sonographic images of thyroid nodules and using the windows “paint” application, we correlate the ACR-TIRADS classification image with the elastosonography image, the anatomy image, the cytological image and the histological image.

Correlations between the sonographic and cytological images were performed by an ultrasound specialist and a pathologist.

This study proposes to investigate the theoretical understanding of pathologies of clinical practice, and according to the Research Ethics Committee of Brazil (CEP), CNS Resolution 510/2016, there is no need for CEP evaluation.

## RESUTS AND DISCUSSION

### Normal Thyroid

#### Anatomy

The thyroid gland consists of two lobes connected to each other by the isthmus and is located in the anterior and central part of the neck below the laryngeal cartilage (10). The thyroid gland is anatomically related to the infrahyoid muscles, namely the sternothyroid, superior belly of the omohyoid and sternohyoid anteriorly, to the larynx, pharynx, trachea, esophagus, external laryngeal and recurrent laryngeal muscles medially, and to the common carotid artery, internal jugular vein and vagus nerve laterally. The normal volume of the thyroid in the adult individual varies between 6.0cm^3^ and 16.0cm^3^, and its weight may vary in the period between 6 and 15 years from 5g to 16g in males, and 5g to 15g in females (11).

#### Ultrasonography

The sonographic image of normal thyroid parenchyma may vary between individuals according to the cellularity or the amount of colloid. However, the normal thyroid gland is usually homogeneous, bright, and with a slightly increased echogenicity relative to the surrounding muscles (12).

#### Elastosonography

Shear wave elastography (SWE) is a new technology that is able of producing complement data relative to tissue elasticity. In elastography the normal values of thyroid tissue in normal individuals are presented in kilopascals (kPa) and the shear wave velocity is expressed in meters per second (m/s) (13).

Studies demonstrated of the elastography values of the normal thyroid gland, and the results ranged from 1.60 ± 0.18 m/s for pSWE to 2.6 ± 1.8 m/s for 2-D SWE (14).

#### Cytology

Fine needle aspiration cytology (FNAC) biopsy is pointed out as gold pattern diagnostic tool in thyroid nodules (15). Benign FNAC findings avoid unnecessary procedures. The FNAC specimens of benign thyroid usually have follicular cells with variable amount of thick and thin colloid (16).

#### Histology

The normal thyroid gland is composed of two basics types of epithelial cells: the follicular cells, which transform iodine into thyroxine and triiodothyronine, and the parafollicular or C-cells, which contain calcitonin (17).

The thyroid gland is involved by a capsule, consisting of dense coating connective tissue and designates septa for thyroid parenchyma, distributed in multiples lobules, and each thyroid lobule includes 20 to 40 follicles. Another group of thyroid secretory cells is of C cells, which secrete calcitonin. The C cells constitutes around 0.1% of the thyroid gland (18).

Figure 3 shows the composition of images that correspond to the normal thyroid (ultrasound, elastography, anatomy, cytology and histology).

**Figure 3.**
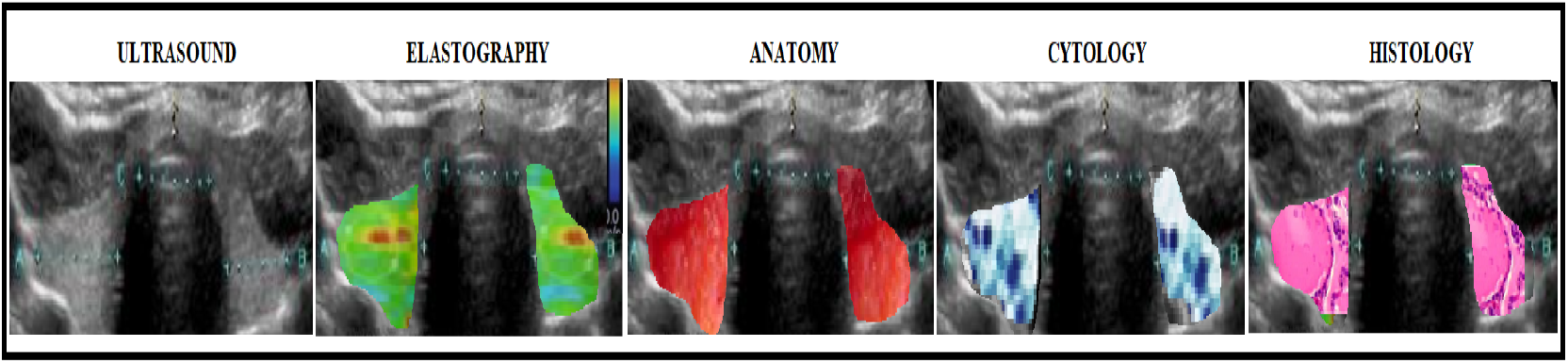
Normal Thyroid. **Source**: Research result

### ACR-TIRADS 1

#### Ultrasonography

After the large-scale use of thyroid ultrasound has been notorious that in euthyroid nodular goitre the most of the nodules is cystic or partially cystic. Composition: Cystic or almost = 0 points, completely cystic or spongiform = 0 points; Echogenicity: Anechoic = 0 points; Shape: Wider-than-tall = 0 points; Margin: Smooth = 0 points, and Echogenic foci: None or comet-tail artifacts = 0 points. Thus, nodules receiving a final score of zero are classified as ACR-TIRADS 1 (5).

#### Elastosonography

Most studies with elastosonography of thyroid have been excluding cystic and/or calcified nodules or predominantly cystic nodules because real-time qualitative thyroid elastosonography only assumes malignancy risk when predominantly cystic nodules are not included (19).

Thus, elastography is inappropriate for predominantly cystic nodules due to artifacts artifacts that lead to unreliable measurements

#### Anatomy

Anatomically the thyroid cyst presents itself with homogeneous cystic component with a clear or yellowish aspect, with defined margins, capsulated and without extrathyroidal invasion and with a fibro-elastic consistency (20).

#### Cytology

Aspirates that contain only cyst fluid are inadequate. The cystic colloid nodules contain only central colloid enclosure with a thin rim of benign follicular epithelium, which justifies the usual non-existence of follicular epithelium in aspirates, and these findings have a low risk of malignancy (21).

#### Histology

The colloid nodular goiter histologically presents follicles of widely varying sizes. Some follicles are much larger than normal and contain abundant colloid. Others resemble normal follicles or are small. The follicular cells may be columnar, flattened, or cubic, indicating variation in functional activity. Beams of fibrous tissue give the gland a nodular appearance (22).

Figure 4 shows the composition of images that correspond to the colloid nodular goiter (ultrasound, elastography, anatomy, cytology and histology).

**Figure 4.**
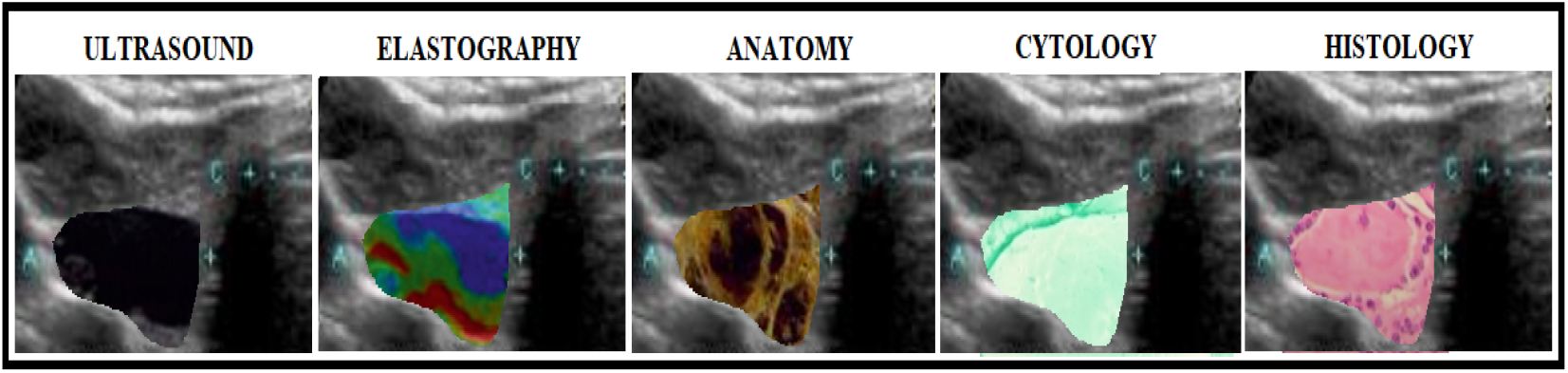
Colloid Nodular Goiter - Benign. **Source**: Research result

### ACR-TIRADS 2

#### Ultrasonography

According to the ACR TI-RADS categorization, the TI-RADS 2 nodule presents the following sonographic characteristics smooth margins = point 0, wider than taller = point 0, hypoechoic = point 2, spongiform composition (0), and with no echogenic foci = point 0. Nodule with capsule, mixed, with solid and hyperechogenic areas hyperechogenic or anechogenic with hyperechogenic spots may show vascularization (5). Anechoic nodule, avascular, and with internal echoes (colloid type 1). Isoechoic nodule with peripheral halo may show hyperechogenic spots (which are not microcalcifications). Does not demonstrate expansibility (no bulging of the thyroid lobe contours) (colloid type 2). Isoechoic nodule with cystic areas may show expansibility (colloid type 3).

#### Elastosonography

In elastography image of a hyperplastic thyroid nodule the largest number of the nodules show relatively not very rigid color elasticity signal. It has been shown that in a ACR TIRADS 2 nodule the mean value of the shear wave are 2.56 ± 0.17 m/s (23).

#### Anatomy

The anatomical specimen of an ACR-TIRADS 2 nodule presents as a nodule usually with a thin fibrous capsule, some contain colloid, and others are hemorrhagic (24).

#### Cytology

Hyperplastic nodules may suffer cystic degeneration and remedial modifications; stromal fibrosis, hemorrhage, hyalinization, calcification, and necrosis can happen. Large colloid pools can form without or with atrophy of epithelium. The aspirates present foamy cells, macrophages filled with hemosiderin, watery colloid, granular protein debris, and scarcity of epithelium and fibroblasts. The reactive cyst enclosure cells possess characteristic cytoplasmic contours, detached to consistent granular cytoplasm, abrupt too much unipolar/ bipolar cytoplasmic processes. Nuclei can be big, rounded to oval but always with regular edges, being able to present grooves and hypochromasia with distinguished nucleoli (25).

#### Histology

In hyperplastic goiter the histology demonstrates a accentuated increase of follicles–a few times huge–and lowering of epithelium. Nodules present a thick viscous substance composed of thyroglobulin. It is rational that these nodules are whether it is the result of a defect of intraluminal thyroglobulin reabsorption (26).

The figure 5 shows the composition of images that correspond to the hyperplastic nodules (ultrasound, elastography, anatomy, cytology and histology).

**Figure 5.**
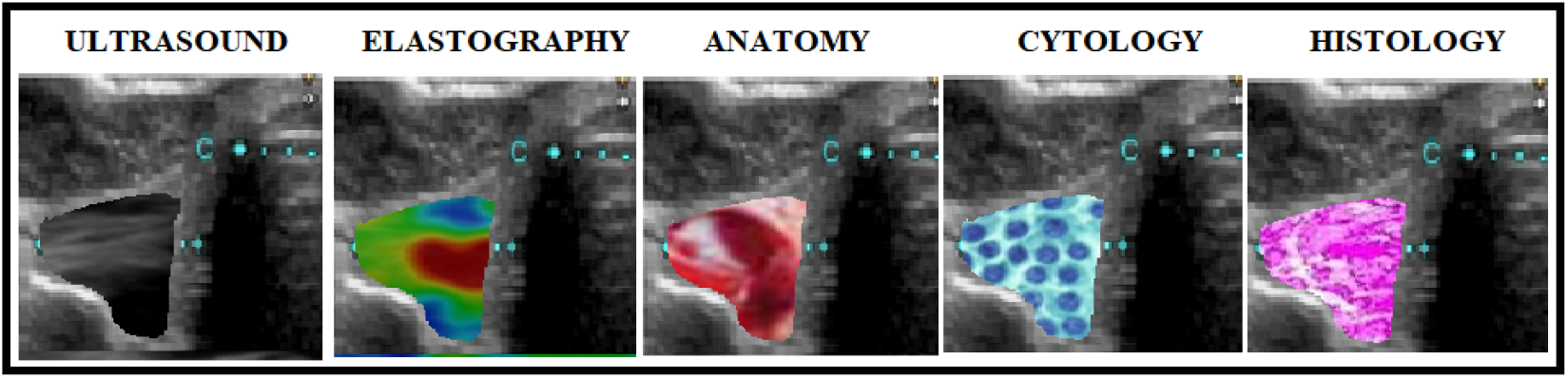
Hyperplastic / adenomatoid nodules thyroid - Benign. **Source**: Research result.

### ACR-TIRADS 3

#### Ultrasonography

Mixed nodule with solid and cystic components, in the case of mixed nodules, only the solid component should be used to score the categories of echogenicity, margins and echogenic foci. Or, completely solid nodule, with echogenicity similar to the rest of the thyroid parenchyma, presenting a hypoechogenic halo that should not be considered for scoring echogenicity or margins. In total there were 3 points, being classified as ACR-TIRADS 3 (5).

#### Elastosonography

The nodule is prominently in green with scarce blue areas with low kPa values and elastographic proportions below than 1, referring that the nodule is softer than the peripheral normal thyroid tissue (27).

#### Anatomy

Anatomically, an ACR-TIRAS 3 nodule are solitary, encapsulated nodules with a thin capsule, usually occur in adults, and are more common in women. Evaluation of the capsule is of fundamental importance in differentiating it from a follicular carcinoma (28).

#### Cytology

The cytological sample contains follicular cells, lymphoid or various cells with atypia or cellular matrix that do not serve the criteria to categorize it in other classification. The most usual possibilities in this category are mentioned in the following circumstances: existence of micro follicles that do not attend the patterns of the hypothesis follicular neoplasm; Hürtle cells predominance in aspirated content with low cellularity and scarce colloid; cytological atypia undermined by pre-analytical artifacts; specimen composed only by Hürtle cells; hegemony of follicular cells of benign appearance, however including focal areas that suggest papillary carcinoma; predominance of follicular cells with benign aspect but containing coating cystic that resemble atypical cells due to the presence of cracks, enlarged nuclei and nucleoli; and finally, atypical lymphoid infiltrate in which the atypia degree is insufficient to categorize it as suspicious for malignancy (29).

#### Histology

Histologically, Bethesda category III consists of small follicles, much smaller than normal and usually without colloid. The tumor is bounded by a fibrous capsule, and the appearance of the neoplastic tissue is regular in all areas. There are no cellular atypia, mitosis, or necrosis. There is no neoplastic infiltration of the capsule or vessels, which is the only reliable criterion for differentiating adenoma from adenocarcinoma (30).

The figure 6 shows the composition of images that correspond to the hyperplastic nodules (ultrasound, elastography, anatomy, cytology and histology).

**Figure 6.**
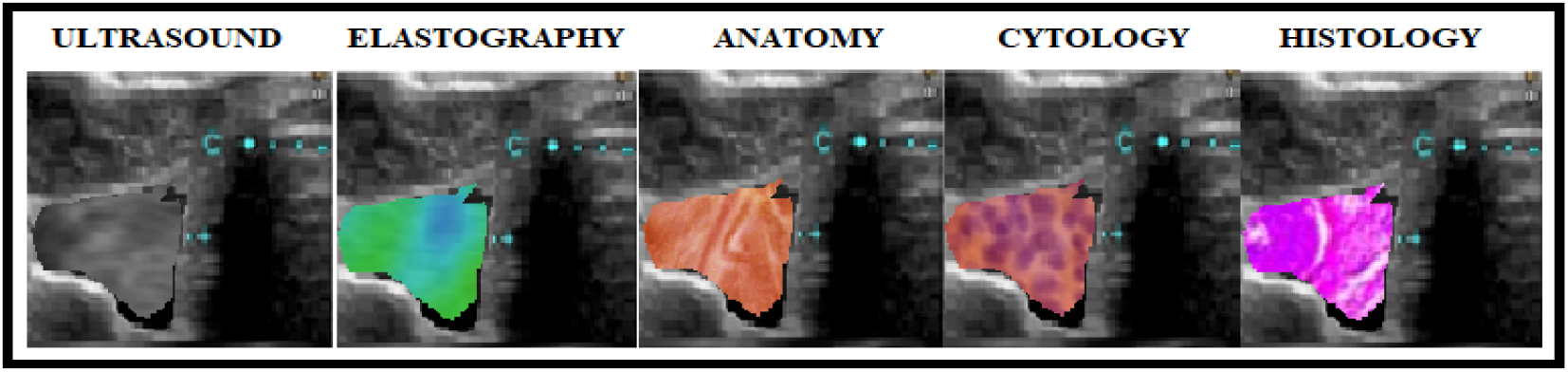
Atypia of Undetermined Significance / Follicular Lesion of Undetermined Significance. **Source**: Research result

### ACR-TIRADS 4

#### Ultrasonography

Solid nodule (2 points), markedly hypoechoic (3 points), wider than high (0 points), smooth margins (0 points), no echogenic foci or posterior attenuation artifacts (0 points). Or a nodule with the following sonographic characteristics, mixed solid-cystic nodule (1 point), isoechoic (1 point), wider than high (0 points), extending beyond the anterior thyroid margin (3 points), without echogenic foci or posterior attenuation artifacts (0 points). In total there were 5 points, being classified as ACR-TIRADS 4 (5).

#### Elastosonography

Nodule presents on elastography as a low elastic nodule, without a rigid border, without internal color homogeneity, no tension in the whole lesion, with blue shading and with focal green dots (31).

#### Anatomy

The macroscopic characteristics of nodules classified as ACR-TIRADS 4, the nodule in most cases, presents as a solid nodule, homogeneous in appearance and pinkish in color, without hemorrhagic or necrotic foci with well-defined margins, with a tenuous capsule, without extra thyroid invasion and of elastic consistency (32). It is not possible to distinguish it macroscopically from a follicular carcinoma. For this, a thorough microscopic study of the tumor capsule is indispensable, to look for infiltration of the capsule itself or of vessels.

#### Cytology

The cytology smear normally has high level cellularity with little or no colloid. It also presents modifications in cytoarchitecture where follicular cells are arranged mostly in microfollicular arrangements and syncytial plaques. Around 16% to 25% of the cases in this category are hyperplastic increases of Hürthle cell in nodular goiter or in a lymphocytic thyroiditis, while 15% to 45% of nodules are malignant (29).

#### Histology

Histologically, follicular adenoma of the thyroid consists of small follicles, much smaller than normal and usually without colloid. The tumor is bounded by a fibrous capsule, and the appearance of the neoplastic tissue is regular in all areas. There are no cellular atypia, mitosis, or necrosis. There is no neoplastic infiltration of the capsule or vessels, which is the only reliable criterion for differentiating adenoma from adenocarcinoma. The thyroid tissue around the tumor shows compression atrophy. Microscopically, follicular carcinoma more or less faithfully reproduces the follicular architecture of the thyroid (24).

The figure 7 shows the composition of images that correspond to the follicular thyroid tumor (ultrasound, elastography, anatomy, cytology and histology).

**Figure 7.**
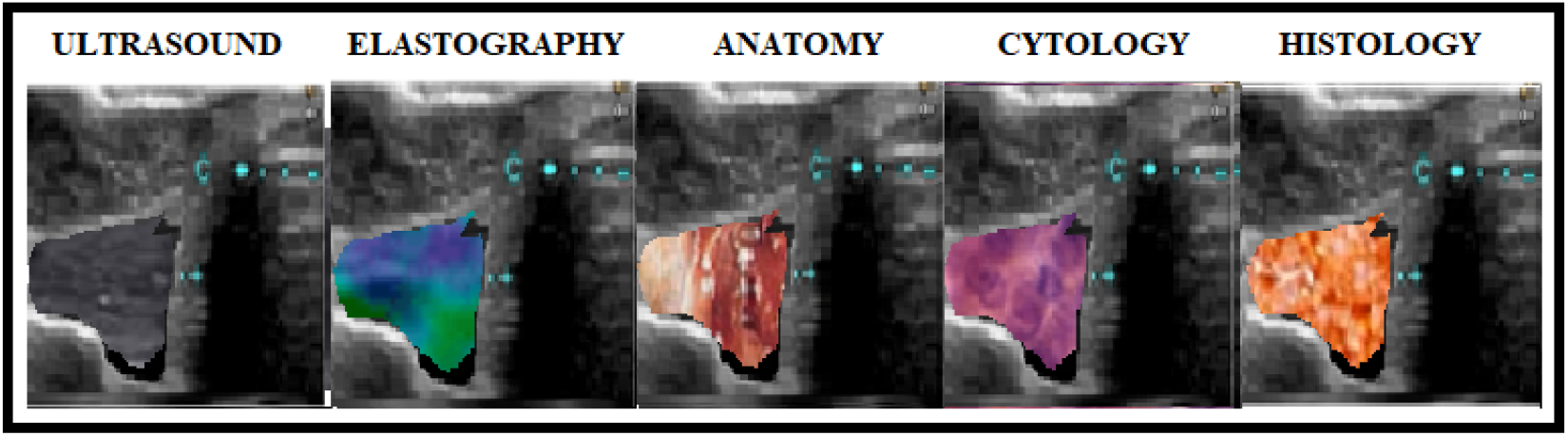
Follicular neoplasm or Suspicious for a Follicular neoplasm. **Source**: Research result

### ACR-TIRADS 5

#### Ultrasonography

Solid nodule (2 points), very hypoechogenic (3 points), lobulated margin (2 points), circumscribed, not parallel to the skin (3 points), presence of peripheral calcifications (2 point) and macrocalcifications (1 point), which corresponds to the ACR-TIRADS 5 classification. Or even, solid nodule (2 points), hypoechogenic (3 points), with irregular margins (2 points) with invasion of adjacent structures (3 points), not parallel to the skin (3 points), also classified as TI-RADS 5. Similarly, a solid nodular image (2 point), hypoechoic (2 points), not parallel to the skin (3 points), regular and circumscribed, totaling 7 points, is classified as ACR-TIRADS 5 (5).

#### Elastosonography

Elastography improves the specificity of gray-scale ultrasonography in characterization malignant lesions. The malignant nodule at elastography is characterized by the lack of elasticity of the tumor tissue, being a harder nodule, defined at elastography by the blue color, when compared to the surrounding tissue of elasticity considered normal. However, the sensitivity of elastography in the recognition of a malignant nodule has a wide variation among the various published studies due to the use of different score systems or different cut-off points as a predictive value of malignancy (33).

#### Anatomy

The macroscopic anatomy of a thyroid carcinoma is described as a solid nodule, whitish and homogeneous in appearance, most often without necrotic foci or hemorrhages, firm in consistency and without capsular invasion (32).

#### Cytology

The papillary carcinoma is the most prevalent cancer in cytological diagnosis among all thyroid cancers. It is the most prevalent diagnosis in thyroid cytology among thyroid cancers. It is identified due to components of thin or thick papillary tissue, presenting fibrovascular nuclei, focal grouping of juxtaposed tumor cells, regular nuclear contours, grooved nuclei and cytoplasmic inclusions within the nuclei. Metaplastic squamous cells and psammomatous bodies can also be found (29).

#### Histology

Histologically, the cells tend to be arranged along branched connective-vascular axes, forming papillae. The cells are usually cuboidal or columnar in shape, uniform and well differentiated. Currently, papillary architecture is no longer considered the decisive criterion for the diagnosis of this type of tumor, and greater importance is given to nuclear features. The chromatin of papillary carcinoma cells is thin and regularly dispersed, giving the nucleus a clear and relatively homogeneous appearance. This is called the ground-glass appearance. There is also a strengthening at the periphery of the nucleus, the chromatin being denser below the nuclear membrane. The intranuclear pseudo inclusions result from invaginations of the nuclear membrane containing cytoplasm. The image is created by superimposition and is not a true inclusion, in the sense of an intranuclear corpuscle. They are a common feature of papillary carcinoma. Faces in nuclear membrane are folds of the membrane that, depending on the plane of observation, give the appearance of a crack in the nucleus, resembling a coffee bean. In some transverse cut nuclei the fold of the nuclear membrane is clearly visible. Psamomatic bodies are calcified corpuscles, often with a concentric lamellated structure, usually found in the connective axons of papillae. They have importance for the diagnosis of papillary carcinoma because they are much rarer in other types of thyroid carcinoma such as follicular, medullary and anaplastic. Also, the folicular component in papillary carcinomas has at least some follicles in the middle of the papillae. In some, the follicular component may greatly predominate over the papillary, or the tumor may be exclusively follicular. In this case, the diagnosis of papillary carcinoma is based on the nuclear features. These papillary carcinomas are called follicular variant and behave biologically like the usual papillary carcinomas, i.e. they are locally infiltrative and usually not encapsulated. Their prognosis is better than that of true follicular carcinoma (24).

The figure 8 shows the composition of images that correspond to the Thyroid Papillary Carcinoma (ultrasound, elastography, anatomy, cytology and histology).

**Figure 8.**
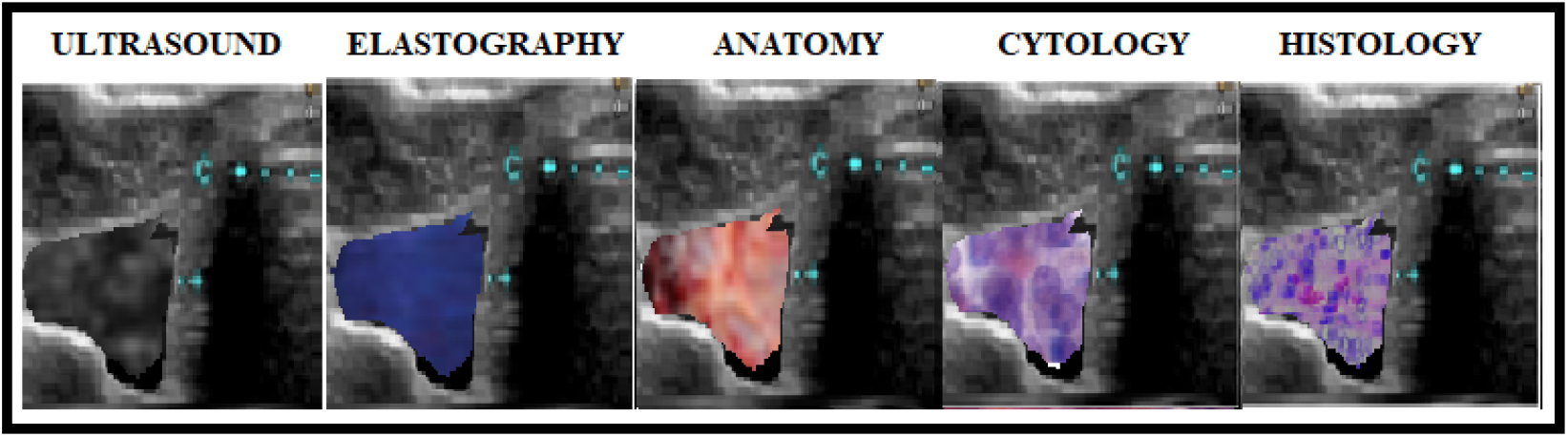
Thyroid Papillary Carcinoma. **Source**: Research result

## CONCLUSION

The sonographic, elastosonographic, anatomical and histological features in the evaluation of thyroid nodules can correlate with the features of the TI-RADS® score, consequently avoiding unnecessary invasive procedures and helping the appropriate clinical/surgical decision making.

